# Leri: a web-server for identifying protein functional networks from evolutionary couplings

**DOI:** 10.1101/2020.12.22.421388

**Authors:** Ngaam J. Cheung, Arun T. John Peter, Benoit Kornmann

## Abstract

Information on the co-evolution of amino acid pairs in a protein can be used for endeavors such as protein engineering, mutation design, and structure prediction. Here we report a method that captures significant determinants of proteins using estimated co-evolution information to identify networks of residues, termed “residue communities”, relevant to protein function. By taking advantage of recent developments in high-performance and parallel computing, we constructed a web-server, *Leri*, that identifies relevant residue communities to allow researchers to investigate how a protein evolves and folds for function(s). All the data of the computational results including high-quality images can be downloaded and presented for publication. This web-server, written in C++, is sufficiently rapid to enable the studies on proteins of up to 400 amino acids.

## Background

Protein sequences, which specify their structures and function, are products of natural selection^1, 2^. The sequences, therefore, harbor evolutionary information. This information in homologous proteins has enabled the accurate computation of residue contacts^3–5^, and helped the prediction of tertiary structures^6–8^. Other useful clues pertaining to protein function might be gleaned from this evolutionary information. With advances in high-throughput sequencing technologies and the expansion of protein databases, we now have insight into evolutionary patterns, which have allowed researchers to detect and identify sets of amino acids performing related functions^9^ and engineer proteins^1^. Statistical coupling analysis (SCA) is a successful approach for identifying sparse networks of interactions among co-variant amino acids (term ‘protein sectors’), for example, the coupled conservation from the networks allows to design of the artificial WW domain (or rsp5-domain) protein sequences that fold into a specified structure representative of the protein family and function as the natural proteins^1, 9, 10^. Moreover, sparse networks of evolutionarily conserved amino acids predict structural motifs for allosteric communication in proteins^11^. This approach is also successful in predicting the effect of mutations on protein function^12–14^. Although it is not difficult to create protein sequences, it remains challenging to design the sequences (being loosely related to natural sequences) that fold in predictable ways and function in a manner indistinguishable from the natural representative of the family. Recent findings have shown that co-evolutionary information hidden in homologous proteins can be harnessed to efficiently create sequences and predict their folding behavior *in vitro*^1, 15^. Thus evolutionary information from homologous sequences provides new insights towards the artificial design of proteins that fold and function in a manner similar to natural proteins.

Biologists have been striving to uncover the mysteries of the relationship between protein primary sequence and their tertiary structures and their function. Despite the vast amount of protein sequence data made publicly available with inexpensive DNA synthesis and next-generation sequencing (NGS)^16–18^, finding biologically meaningful information in the data requires elucidating the evolutionary determinants that give rise to protein functions. These determinants can be manifested as amino acid coevolutionary patterns. Such coevolutionary patterns are informative in an array of applications from protein structure prediction (PSP) constrained by residue contacts (derived from direct information^3, 6^) to protein engineering and drug discovery. Our understanding of protein biology is limited to homologous proteins that preserve similar structural information (e.g., contacts between residues), but challenges remain what and how couplings drive protein to fold and function, specially, how the strongly coupled residues that are sparse and organized into highly ordered architectures link to protein’s functional sites. Technological advances over the years have enabled certain estimations of residues ranked by their importance relevant to protein function, including the ability to infer protein sectors, groups of amino acids that are coevolving in homologous proteins^9^, and assess mutation effects^13^. Assessing the network-level effects (e.g. in partitioned blocks) of evolutionary couplings between pairwise amino acids, however, remains a challenge. Even if experimental investigations (e.g. directed evolution^19^) could provide all components necessary for understanding protein function, discovering interactions among amino acids requires a computational approach to analyze large-scale sequence data and put pieces of information together. The energy changes (the energy difference between the wild-type and mutated sequences, Δ*E* = *E*_*wt*_ *−E*_*mutant*_) computed from the positionally conserved couplings have a strong correlation with the transition temperatures for the extant and ancestral Thioredoxin (Trx) proteins, and those changes might allow engineering proteins with improved properties^20^.

Here, we present an enhanced computational web-server, termed *Leri* (learning engine to recognize life), that allows researchers to detect co-evolution patterns (highly ordered networks of coupled amino acids, community structures in graph theory, termed residue communities). It can also identify functionally important residues that would be used for protein design and folding from either experimental or natural evolution^8, 21^. This provides a systematic method by applying the spectrum analysis on evolutionary coupling analysis (SAEC) for inferring functional networks in proteins, identifying mutations as a guide for protein engineering and facilitating researchers to understand function. Through two instructive examples, we showcase the capabilities of *Leri* in building the residue communities that are relevant to protein’s functions. In the first case, we apply *Leri* to identify the functionally relevant residues in the mitochondrial morphology maintenance 1 (Mmm1) protein. In the second example, we demonstrate how *Leri*’s efficiency in capturing the residues involved in lipid transportation in the endoplasmic reticulum-mitochondria encounter structure (ERMES) complex.

## Results

### Implementations

Leri is written in C++ and provides visualizations of the results in HTML format. Plotly^22^ is employed to visualize the data interactive web graphics, and NGL Viewer^23^ is used to process and view protein structure files.

The web implementation of *Leri* allows users to choose an engine for a specific computation on a given protein sequence, as illustrated in Fig. 1. The *Leri* web-server provides four types of computations that are all driven by a query sequence, including: (1) evolutionary coupling analysis and functional networks to capture evolutionary signatures (e.g. residue communities) and residues evolved for function; (2) protein single mutation for evaluating mutants for potential applications in protein engineering; (3) protein coupled mutations result in multiple mutants that have evolutionary dependence on each other; and (4) protein sequence design from the inferred residue communities can guide the engineering of functional proteins with altered (bio)chemical activities. Broadly, to create analytical results of a protein of interest, the user needs to provides its sequence or multiple sequence alignment (MSA) in FASTA format, together with an optional PDB file, on which the identified “residue communities” are mapped, and all the residue communities are computed from the MSA of each protein by the SAEC method (see Methods).

**Figure 1.**
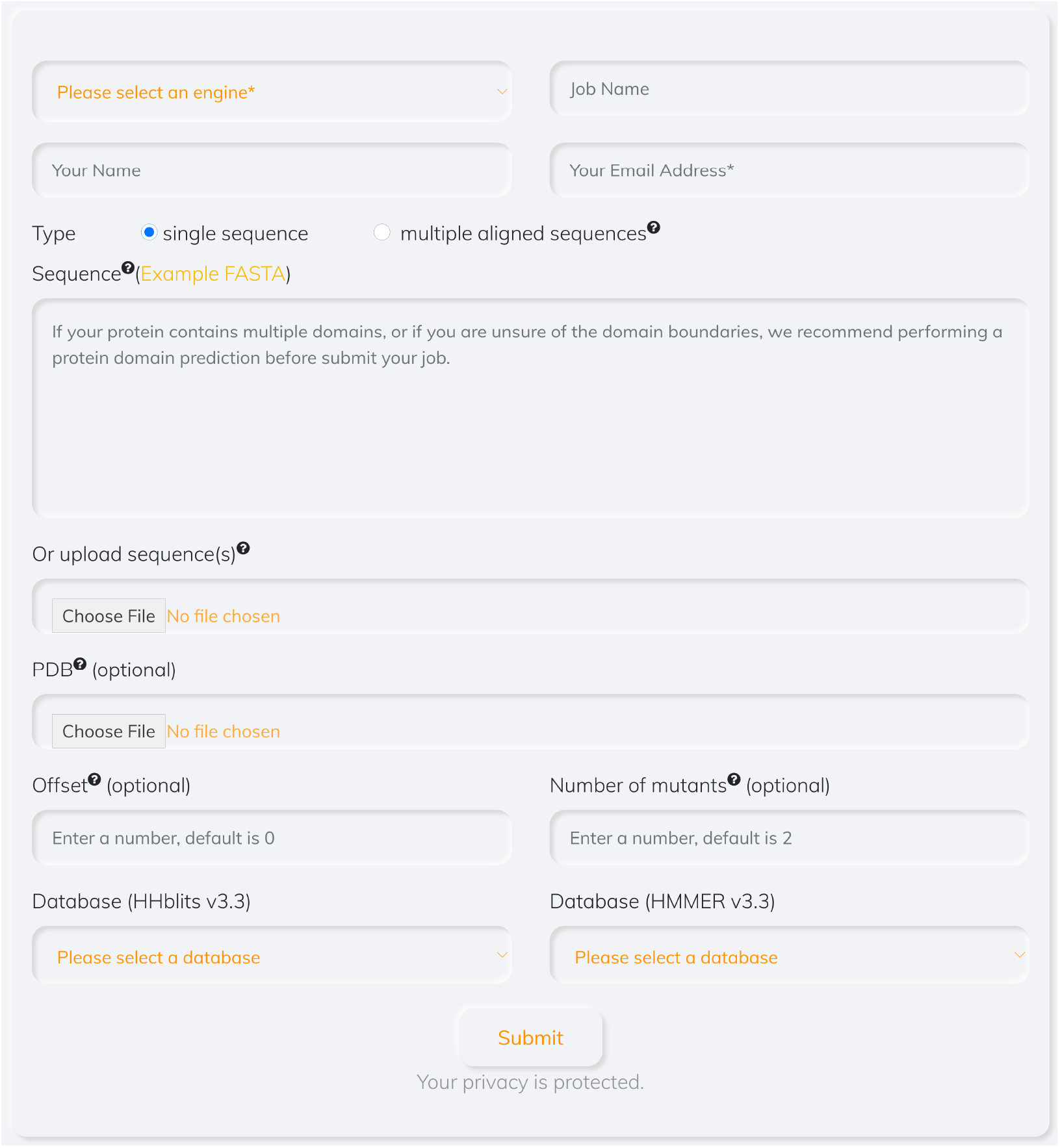
Protein sequence submission page for the Leri web server. Users are prompted to select a computing engine and upload/paste a sequence in FASTA format to submit for calculations.

On completion of calculations, a report in HTML format is presented with a summary of detailed results by interactive graphics and visualization of the inferred residue communities in the NGL Viewer^23^ (if protein structure is provided) (Fig. 2). The output data (as CSV and plain text files) can be downloaded for customized use. Downloads are also available for the 3D representation (as PyMOL script) of the structure with mapped residue communities. A summary is given for the basic information of the job (Fig. 2a). As detailed, the MSA can be visualized for either an uploaded sequence alignment or an alignment generated against the database, while the distribution of sequence identities between pairwise sequences are interactively shown in histogram chart (Fig. 2a and b). To measure the information on residue conservation, the relative entropy (Kullback–Leibler divergence^26^) is computed for the most prevalent residue at the specific position (Fig. 2d). The entropy evaluates how different the observed amino acid A at the *i*th position would be if A randomly occurred with an expected probability distribution. The user can measure interactions among amino acids based on the evolutionary coupling strength (Fig 2e) for a particular protein and infer its interaction/binding sites that are estimated in the residue communities (Fig. 2f-h). The amino acids located in those communities are functionally important and are more sensitive to substitutions, whereas the residue outside of the communities are tolerant to mutation (Fig. 2i). The differences between the wild-type and an optimized mutant sequences (with the lowest sequence energy in a fixed number of iterations) are computed from any pairwise mutants (Fig. 2j), and the predicted energy differences have been demonstrated to be relevant to the thermostatically stability of proteins^20^. On the web server, the number of coupled mutants can be customized, and the energy can be a measurement to guide the design of a protein of interest.

**Figure 2.**
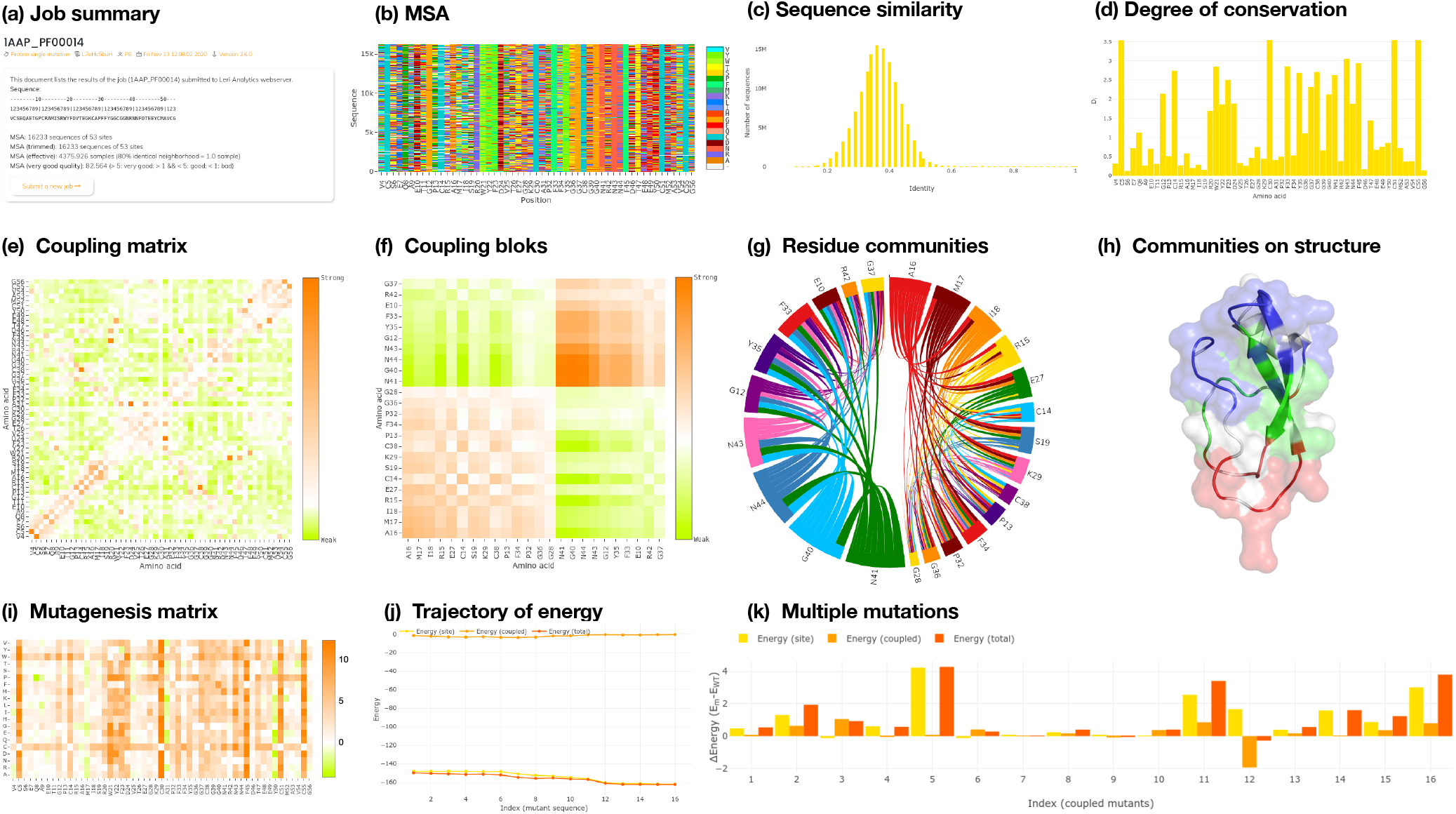
Interactive results shown in HTML format for an Leri web-server job. (a) Basic summary of the job submitted by a user. (b) The MSA is collected from Pfam database^24^ (PF00014). (c) Sequence similarity. (d) Degree of conservation at the position of each amino acid. (e) evolutionary coupling matrix inferred from the MSA by the ECA method. (f) and (g) show the top two residue communities that are computed from the coupling matrix. Interactions with both positive and negative values are illustrated in (f), while positive interactions are shown in an interactive chord graphic in (g). (h) Mapping the inferred residue communities to the tertiary structure (PDB: 1AAP) in red, blue and green in interactive web interface based on NGL Viewer^23^, PyMOL^25^ script can be downloaded for local generation. (i) Substitution matrix for 20 common amino acids on the WT protein sequence. (j) Energy trajectory of the mutant sequence starting from the WT sequence. (k) Energy differences of coupled mutants between the WT and mutant sequences.

To draw interpretable conclusions from the results, we provide interactive visualizations and quantitative measurements (e.g. energies) of the data on the *Leri* webpages. In Fig. 2g, for example, we present an interactive circular plot of the top two residue communities that consist of residues in highly ordered patterns (Fig. 2f) with coupling strength between them. The communities can be also mapped on the tertiary structure (Fig. 2h) if provided, and structurally visualized for the dominant signatures via the inferences of the SAEC method.

### Phospholipid-binding site inferred by coupling networks

The endoplasmic reticulum-mitochondria encounter structure (ERMES) complex, consisting of at least four proteins^27^: Mdm10, which is integral outer mitochondrial membrane (OMM) protein, Mmm1, which is integral to the ER membrane, and Mdm12 and Mdm34, which are a cytosolic proteins, have been functionally connected to phospholipid biosynthesis^27^. Mmm1 binds phospholipid molecules^28^ (Fig. 3b) through a conserved domain called SMP (Synaptotagmin-like, Mitochondrial and lipid-binding Proteins) that also present in Mdm12 and Mdm34. This first case illustrates how the inferred communities of residues can be used to identify functional sites in Mmm1 protein. The couplings between pairwise amino acids were inferred from the alignment of natural evolution sequences. The highly coupled residues fall within three networks (Fig. S1), and the top two networks consist of residues that have distinct intra-interactions (Fig. 3a) and are highlighted in red and blue in Fig. 3b, while the other network (green in Fig. 3b) includes residues that have inter-interactions with those in other two networks.

**Figure 3.**
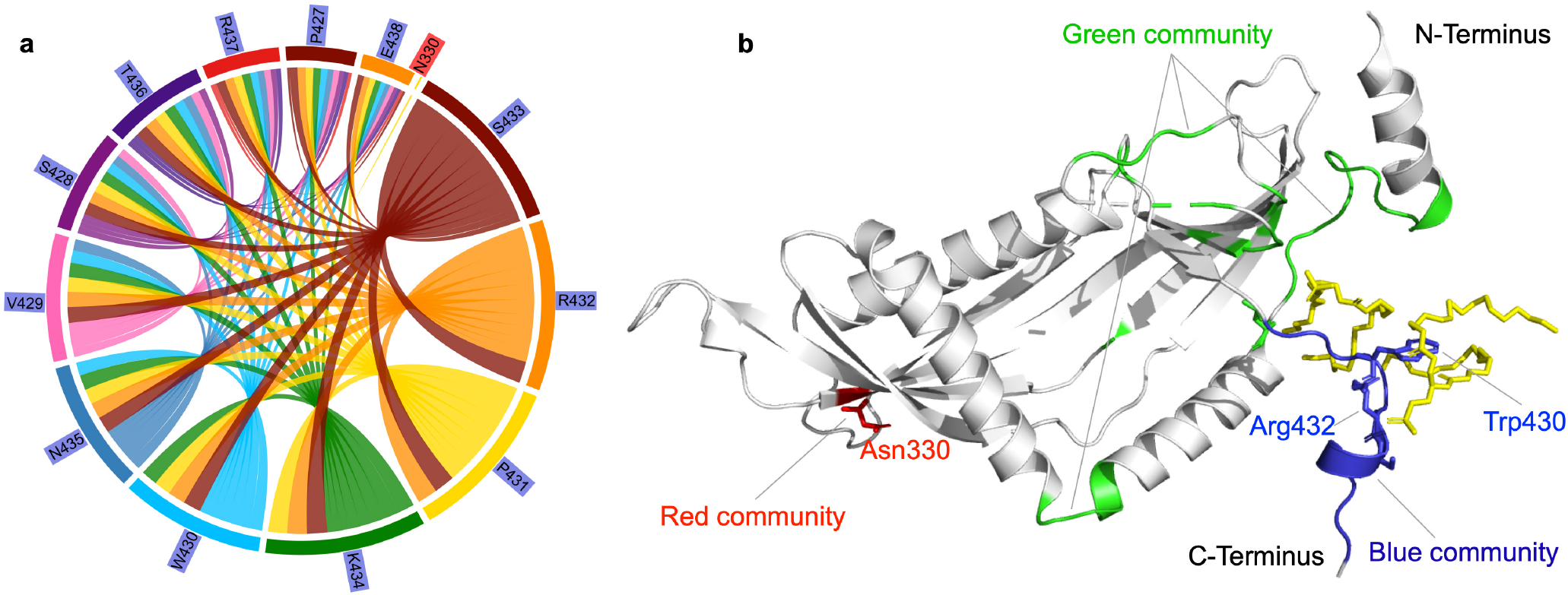
Function-related amino acids of Mmm1 protein (PDB ID: 5YK6) are identified by the residue communities. (a) Coupling between pairwise amino acids in the top two residue communities (the residue N330 is the only one in the red community). (b) The top three residue communities are mapped to the tertiary structure. Residues in three communities are shown in red, green, and blue, respectively.

The residues in the well-ordered loop (residues Y208-S218) are important to mediate the self-association of the Mmm1 dimer and phospholipid-binding at the highly conserved C-terminus^29^. As illustrated in Fig. 3b, those residues are captured from the MSA of the Mmm1 using the proposed SAEC method. That is, the inferences are consistent with experimental results. Interestingly, we find that the residues positioned at the blue communities engage in the binding specificity of phospholipid through the conserved Trp430 and Arg432 residues (Fig. 2b) that are demonstrated to be biologically essential^29^.

### Communities of coupled residues in the Mmm1-Mdm12 complex

As a further way to explore the inferences, we assessed whether residue communities play a role in protein-protein interaction and lipid transport. In practice, we performed calculations on the Mmm1-Mdm12 complex to identify the determinants in those communities. To achieve accurate inferences, the MSA of each domain in the complex are searched against the sequence profile database (see Methods). The estimated residue communities of each domain are mapped to the structure of the Mmm1-Mdm12 complex (Fig. 4).

**Figure 4.**
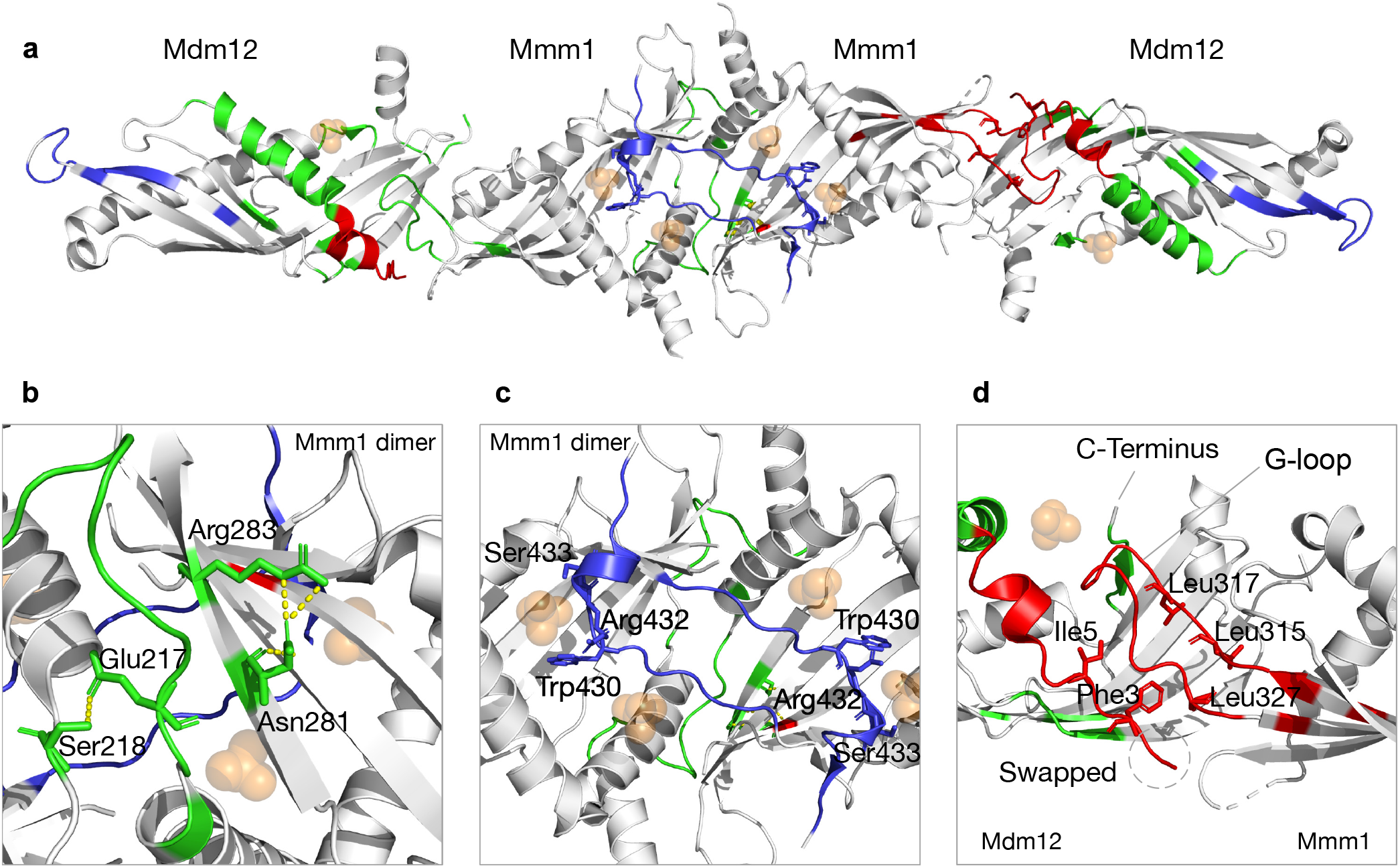
Communities of coupled residues mapping to the tertiary structure of Mmm1-Mdm12 complex (PDB ID: 5YK7). (a) The top three communities of coupled residues are mapped in red, green, and blue. (b) A pattern of co-evolution and the co-evolved residues inferred by the SAEC method are shown in green as stick with polar interaction in yellow dot lines. (c) The inferred residue community highlighted in blue consists of function-related residues. The two highly conserved residues Trp430 and Arg432 are shown as stick in blue. (d) Critical determinants of the interaction between Mmm1 and Mdm12.

From the overall calculations on sequence profiles, we observed distinct residue communities and determinants in the complex of Mmm1-Mdm12 (Fig. 4a). As investigated^29–31^, Mmm1 interacts with Mdm34 (Fig. S2) through Mdm12 via relatively weak or transient interactions. In addition, The studies^29–31^ suggested that Mdm34 and Mdm12 interact with each other through their N termini, specially via *β* -strand swapping^31^ (residues in blue at the N-terminus of the Mdm12 as shown Fig. 4d). The identified residues between the Mmm1 and Mdm12 suggest the three proteins, Mmm1, Mdm12, and Mdm34, can bind to each other at the interaction sites within the predicted residue communities (Fig. 4a). The binding interface between Mmm1 monomers is characterized by the blue and green communities, and the green community consists of a well-ordered loop that contacts the head region of the other molecule of the Mmm1 dimer. Moreover, the G-loop, as shown in blue community, (the extended hairpin loop between at C-terminus^31^) of the Mmm1 monomer plugs into Mdm12, and it introduces conformational change when the complex assembles^29^ (Fig. 5). The co-evolution is distinctly revealed by two residues, Asn281 and Arg283 (Fig. 4b), among Mmm1 orthologs.

**Figure 5.**
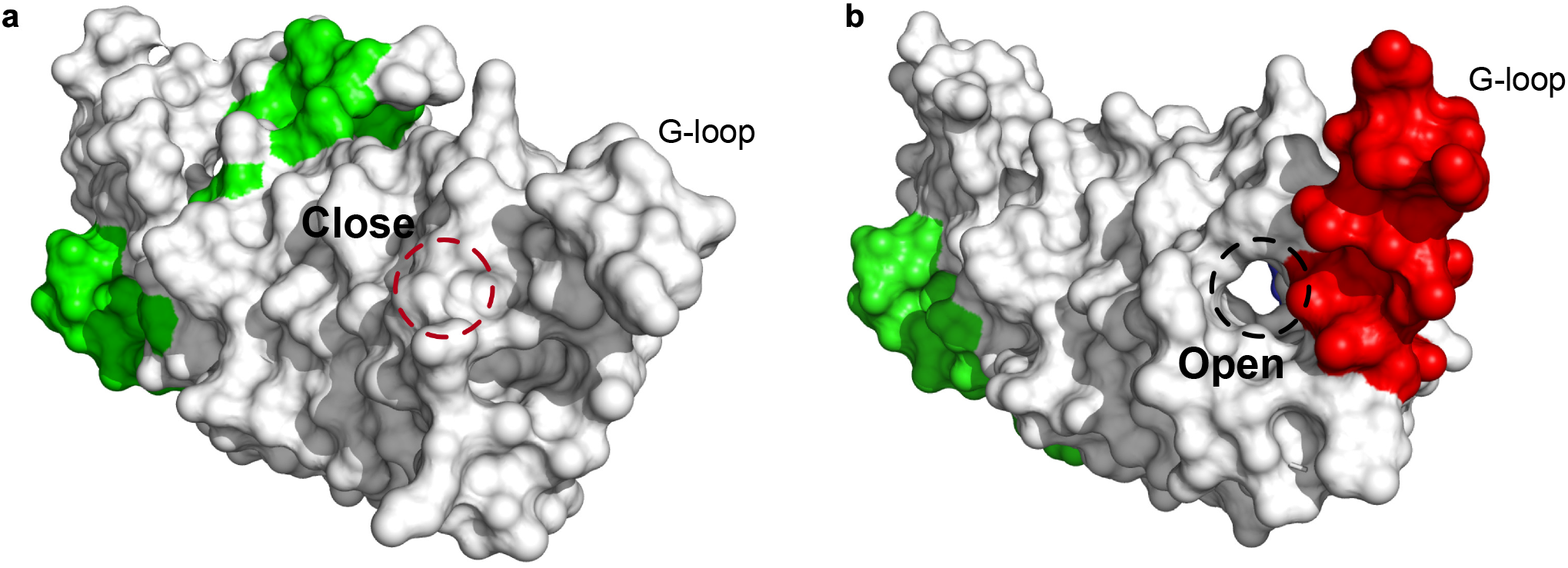
The G-loop of Mmm1. The inferred residue communities are highlighted in green and red, and the dashed circles in red and black are to indicted the position of the channel (open or close). (a) The G-loop in a single domain (PDB: 5YK6) is not captured by the SAEC method. (b) The G-loop causing conformational change in the Mmm1-Mdm12 complex (PDB: 5YK7) is identified from the homologous sequences.

The G-loop (consisting of residues 425-432, Fig. 4c) of the Mmm1 covers the concave surface at the center of the dimeric SMP domain, and in the loop, there are two absolutely conserved Trp430 and Arg432 residues that are essential for the recognition of phospholipids. An extensive hydrogen-bonding network (consisting of the conserved Arg253, Arg415, Trp411, Trp430, Arg432, and Ser433) coordinates the phosphate group and carboxyl oxygen of the distal phospholipid. Among those residues, three residues (Trp430, Arg432, and Ser433) are statistically significant in the residue communities that are inferred by the SAEC method, which is consistent with their contribution to homodimerization for the lipid coordination in Mmm1. The G-loop of Mmm1 can plug into the head region of Mdm12 and cover the solvent-exposed concave surface of Mdm12 in an extended conformation (Fig. 4d and Fig. 5). In particular, the hydrophobic amino acids Leu315, Leu317, and Leu327 in Mmm1 form extensive and coordinate nonpolar contacts with the side chains of Phe3, and Ile5 of Mdm12 (Fig. 4d). Those hydrophobic contacts play a critical role in the Mdm12-Mmm1 interaction.

A conformational change of the G-loop make an extended structure to plug into the head region of the scMdm12*δ* and cover the solvent-exposed concave surface of scMdm12*δ* ^29^. The spectrum of the couplings among amino acids is consistent with the conformational changes of the G-loop as illustrated in Fig. 5. To demonstrate that residue community (in red) is relevant to the conformational change, the two structures of Mmm1 as single domain and in the complex are aligned to each other for comparison (Fig. 5). The statistical qualities of the residue communities (in green) do not capture the important information from the G-loop when Mmm1 adopts a shape as a monomer, while the Mmm1 changes its shape via the G-loop (residue community in red) in response to the formation of a complex between the Mmm1 and Mdm12 proteins.

## Materials and methods

### Framework and implementation

*The leri* web-server takes a single sequence or a multiple sequence alignment (MSA) as input to infer evolutionary couplings and residue communities that could be determinants of functional specificity. Given an MSA (Fig. 6a, whatever provided by the user or search the target sequence against the databases), quality control, including sequence trimming and re-weighting, is required to improve the estimations of evolutionary information and reduce the background noises. Inferences (site-independent biases and couplings, Fig. 6b) are computed from the controlled MSA, and the co-evolution between pairwise residues is measured by the strength of couplings (Fig. 6c). The residues are identified as determinants of functional specificity (Fig. 6d), and those significant coupled residues are mapped to the tertiary structure of the target sequence if provided (Fig. 6e).

**Figure 6.**
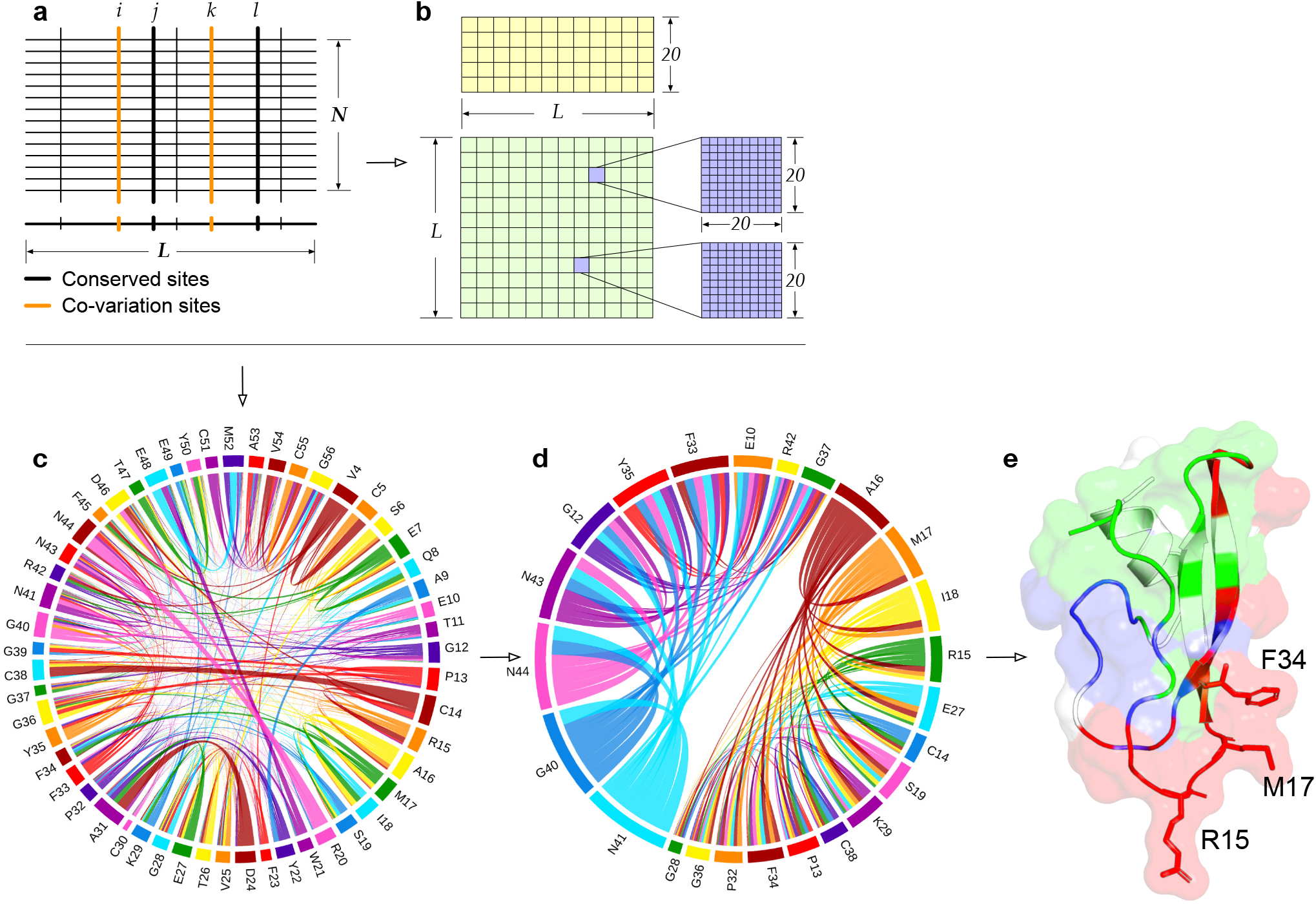
Identification of function-relevant residue communities from a sequence alignment. (a) Conserved and co-varied residues in an MSA. (b) Energy-like potentials including site preferences and pairwise couplings are inferred by the SAEC method. (c) Interactions between pairwise residues. (d) The top two residue communities are computed from the pairwise couplings by spectral analysis. (e) The top three communities including a network with inter-interactions between the top two are mapped to the tertiary structure (PDB: 1AAP).

### Generation of multiple sequence alignments

Co-variation patterns in a natural protein family highly depend on the quality of its MSA when the statistical transformation is made from the genetic record. For each analyzed protein, its multiple sequence alignment was obtained from two databases by the profiles HMM (hmmsearch version 3.3)^32^ and HHblits (version 3.3.0)^33^ homology search tools. Firstly, an alignment of a target protein is obtained by searching its sequence against the Uniclust30 database (as of 2/2020) using HHblits^33^ with the default parameters at an E-value threshold 0.001. The other alignment of the same protein is generated by the default five search iterations of *jackhmmer* in the HMM suite (version 3.3)^32^, searching the query sequence against the UniRef90 database (release 3/2019)^34^. Finally, the two alignments were combined and aligned according to the target sequence. The final alignment of each protein was combined from the two obtained alignments according to the query sequence. Thereafter, it was trimmed based on minimum coverage, which satisfies two basic rules^20, 35^: (1) a single site with more than 90% gaps across the MSA will be removed; and (2) a sequence with the percentage of gaps less than a given threshold (80%) will be deleted from the MSA. In the alignment of each protein, the weight, *ω*(*τ*), of each sequence was computed by the sequence identity *I*, which is measured by the normalized Hamming distance *D*_*H*_(*τ, τ*_*j*_) between the sequence *τ* and all other sequences. Accordingly, the weight is defined^13^ as Eq(1).

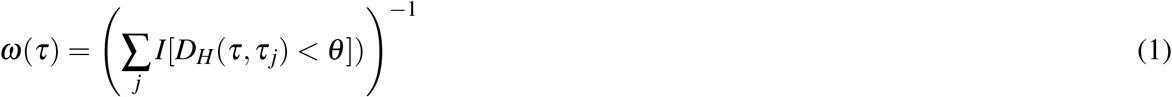

where *θ* is a threshold that controls the maximum diversity of pairwise sequences, and *θ* = 0.2 is default, that is, the threshold of sequence identity is 80% between the two sequences.

### Inference of residue communities

Here, we applied a global probabilistic model^13^ with the pseudo-likelihood maximization approach to capture evolutionary information from the multiple sequence alignment (Fig. 6). A protein sequence *τ* is derived by a probability *P*(*τ*) from a distribution over the space of all possible sequences in its family. The probability *P*(*τ*) is defined as

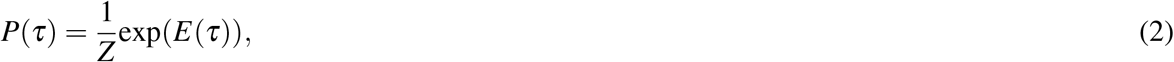

where *Z* is the partition function that normalizes the distribution by summing over the Boltzmann factors of all possible sequences in the protein family. In the model, the measurement of the Boltzmann factors for each sequence *τ* is defined by the evolutionary statistical energy (a Markov Random field or a Potts model in statistical physics) as Eq. (3).

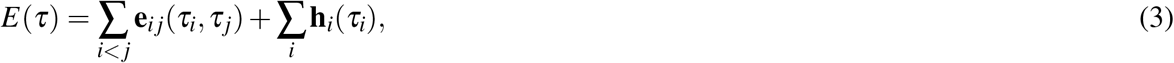

where **h**_*i*_ and **e**_*ij*_ are, respectively, site-specific bias terms and coupling terms between pairwise amino acids.

Our method focuses on discovering intra-interacted residues that underlie distinct residue communities (patterns of co-evolved amino acids, Fig. 6d) within a protein or complex. As the coupling matrix **e**_*ij*_ (Fig. 6b) is to measure the strength between pairwise amino acids at positions *i* and *j* in the protein sequence alignment, it is straightforward to identify significantly coupled pairs by spectral decomposition, which diagonalizes the coupling matrix by linearly combining the positions of the amino acids into eigenmodes. The spectral decomposition provides a way to extract nonrandom correlated modes of the couplings, effectively reduces noises in the couplings, and partially sorts out the residues into different “residue communities”. A procedure of Jacobi’s iteration is carried out to determine the positive eigenvalues and the corresponding eigenvectors. Accordingly, the eigenvalues of the coupling matrix are to indicate the information in the inferred residue communities, while the corresponding eigenvectors give the weights for the positions of amino acids. In order to make the residue communities as much statistically independent as possible, the top three residue communities are defined based on two of the top five eigenvalues and their corresponding eigenvectors (**v**_*k*|*k*=1,…,5_) of the matrix **e**_*ij*_ (Eq 3) as: (1) community I (red) consists of residues at the *i*th position of 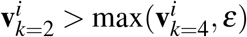; (2) community II (blue) includes residues at the *i*th position of 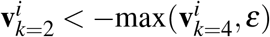; and (3) community III (green) includes residues at the *i*th position of 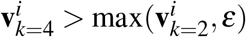. We used an *ε* = 0.05 as the threshold to project the amino acids reduced from the coupling matrix and extract meaningful residue communities.

## Discussion

*Leri* allows fast, systematic, and optimized estimations of the partitioned interactions among amino acids (residue communities) that may not be predictable from physically contacted residues (e.g., the distance between C_*β*_ −C_*β*_ of pairwise residues), such as co-evolving amino acids at function sites/interfaces, and formation of function interface from a network of residue interactions. These inferences can, in turn, provide mechanistic explanations for experimental observations by identifying the key determinants involved in protein function. Accordingly, *Leri* provides a computational framework in which predictions and experimental evidence can be integrated to infer the key principles at the network-level from homologous protein sequences.

In this study, we demonstrated that coupling between amino acid pairs can be grouped into networks, termed residue communities, in which the amino acids have stronger interactions in intra-networks but weaker interactions in inter-networks. We also demonstrate that identified residue communities are functionally relevant in protein specificity, e.g protein-protein interactions, phospholipid-binding activity, and conformational changes.

Although the inferences of the SAEC method are able to predict key residues related to protein function in the highly ordered communities, it might not be successful in identifying the communities if the multiple aligned homologous sequences suffer from limitations such as lack of diversity in the sequences and/or biases arising from evolutionary constraints across a protein family. Moreover, there is still a challenge that remains in computational resources, and this may make it difficult in exploring the whole space of a large protein and interpreting the signatures that are identified in the communities. In spite of those challenges, the proposed SAEC approach, supported by the *Leri* webserver, is readily applicable for analysis and identification of the residues communities that are relevant to protein functions. We anticipate that the analyses from the SAEC method will benefit the scientific community of biochemists and biophysicists for a better understanding of proteins and their subsequent engineering.

## Data and code availability

The web-server is freely accessible at https://kornmann.bioch.ox.ac.uk/leri for non-commercial use. The standalone package of Leri working on Linux system is available upon request for non-commercial use.

## Acknowledgements

We acknowledge Sabine van Schie and Dr. Christian Covill-Cooke for discussion and proofreading the manuscript. We also thank all members in the Kornmann’s Group for helpful discussions. ATJP is supported by the Spark grant of the Swiss National Science Foundation (SNSF).

