## Supplemental Figure for "Leri: a web-server for identifying protein functional networks from evolutionary couplings"

### Supplementary information

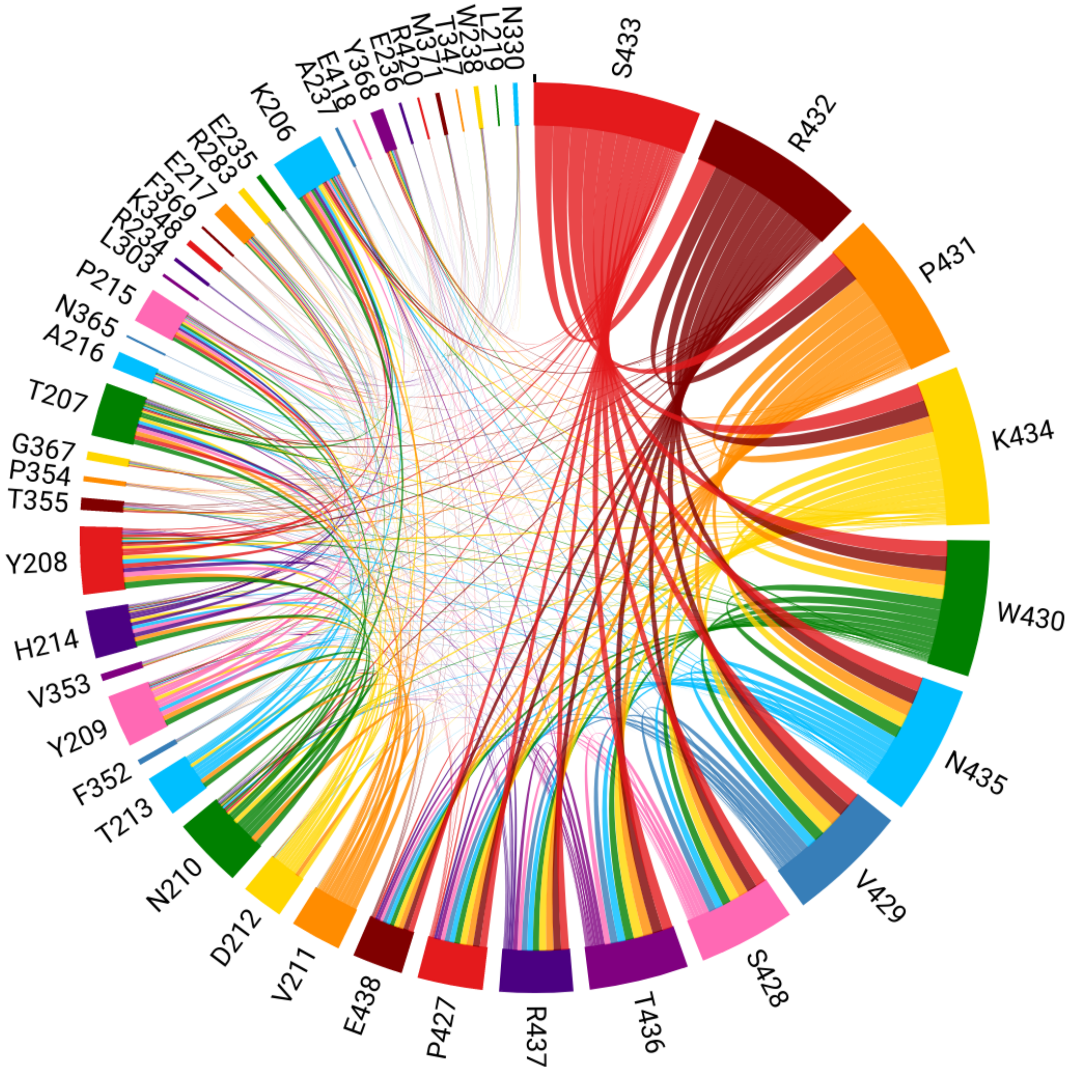


**Figure S1.** Chord plotting of the top three networks inferred from the sequence profiles of the Mmm1.

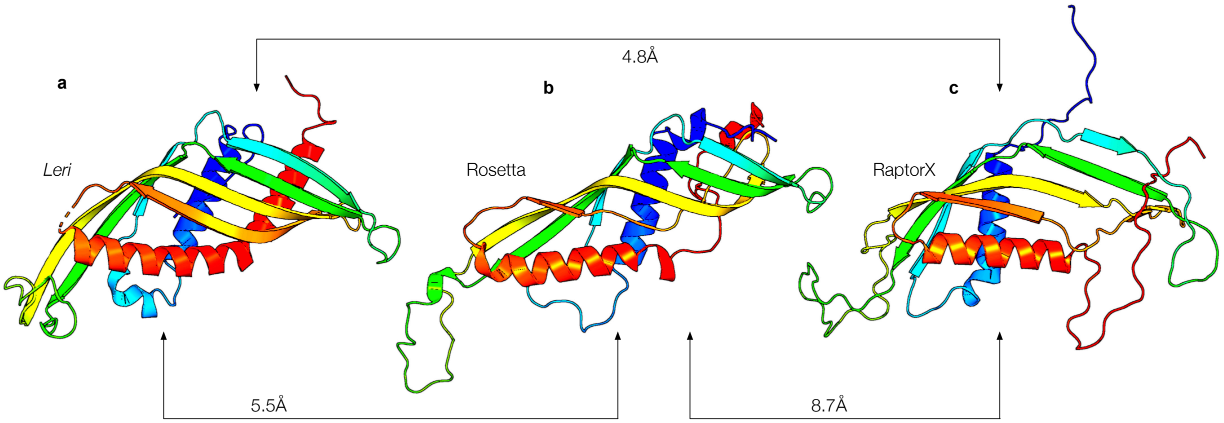


**Figure S2.** The predicted structure of the Mdm34 from the coupled interactions by *Leri* folding module. The predicted structure (best-so-far) is compared to those of Rosetta[?] and RaptorX[?]. The root mean square deviations (RMSDs) are computed by PyMOL[?].
